# Cortico-cortical feedback from V2 exerts a powerful influence over the visually evoked local field potential and associated spike timing in V1

**DOI:** 10.1101/792010

**Authors:** Till S. Hartmann, Sruti Raja, Stephen G. Lomber, Richard T. Born

**Affiliations:** Department of Neurobiology, Harvard Medical School, Boston, MA; Department of Physiology, McGill University, Montréal, Québec, Canada; Perceptive Automata, Somerville, MA

**Keywords:** synchronization, oscillations, rhythm, gamma band, feedback connections, visual cortex, vision

## Abstract

The local field potential (LFP) is generally thought to be dominated by synaptic activity within a few hundred microns of the recording electrode. The sudden onset of a visual stimulus causes a large downward deflection of the LFP recorded in primary visual cortex, known as a visually evoked potential (VEP), followed by rhythmic oscillations in the gamma range (30-80 Hz) that are often in phase with action potentials of nearby neurons. By inactivating higher visual areas that send feedback projections to V1, we produced a large decrease in amplitude of the VEP, and a strong attenuation of gamma rhythms in both the LFP and multi-unit activity, despite an overall increase in neuronal spike rates. Our results argue that much of the recurrent, rhythmic activity measured in V1 is strongly gated by feed-back from higher areas, consistent with models of coincidence detection that result in burst firing by layer 5 pyramidal neurons.

## 1. Introduction

Despite decades of study, the local field potential (LFP) and its associated rhythmic components remain a mysterious and even contentious set of phenomena. While many groups agree that the LFP consists mainly of the summed synaptic activity near the recording electrode (Buzsáki and Wang, 2012; Lindén et al., 2011; Mitzdorf, 1985), there is disagreement as to the extent to which action potentials and other non-synaptic events also contribute (see, *e.g.*, Reimann et al. (2013)) and even to how “local” these sources are (Kajikawa and Schroeder, 2011).

Perhaps even more controversial are LFP oscillations in the gamma band (30-80 Hz), which are reported by many, but whose functional interpretations are agreed upon by few. While some investigators have made the case for a computational role in sensory coding (Buzsáki and Chrobak, 1995; Fries, 2009; Singer, 1999), others have argued that they are simply an epiphenomenon produced by local interactions between excitation and inhibition (Ray and Maunsell, 2015). Several groups have provided evidence indicating that gamma oscillations are a signature of feedforward processing, whereas slower rhythms, like alpha (5-15 Hz) or beta (10-20 Hz), are a marker of feedback signals (Bastos et al., 2015; van Kerkoerle et al., 2014). Regardless of their function, gamma rhythms are believed to be generated locally through strong excitation-inhibition interactions, even though they may become synchronized across relatively large regions of cerebral cortex (Börgers and Kopell, 2003; Buzsáki and Wang, 2012; Headley and Paré, 2017; Tiesinga and Sejnowski, 2009).

Here we examined the influence of cortico-cortical feedback on the local field potential (LFP) and on spiking (multi-unit activity, MUA) by reversibly inactivating retinotopically corresponding parts of V2 and V3 while recording visually evoked activity in V1 of alert, fixating macaque monkeys. Inactivating feedback produced profound effects on the LFP recorded in V1: a large decrease in the visually evoked potential (VEP) after the initial stimulus induced deflection, and a significant reduction of rhythms in the theta-, alpha- and gamma-bands. During control conditions, the MUA oscillated at gamma-band frequencies, and this was completely abolished during feedback inactivation even though visually evoked spike rates were slightly increased. Our results reveal a strong influence of corticocortical feedback on rhythms previously believed to be of purely local origin.

## 2. Results

While two monkeys performed a fixation task for juice rewards, we displayed large (3.6 – 5° in diameter), high-contrast, sinusoidal gratings of varying orientations (Fig. 1d). Such stimuli have been widely found to evoke strong oscillations in the low gamma band (30 to 55 Hz) in early visual areas (Jia et al., 2013b; Ray and Maunsell, 2010), which we confirmed by recording LFPs and MUA with multi-electrode arrays implanted in V1 (Fig. 1a,b). We then reversibly inactivated retinotopically aligned parts of V2 (Fig. 1c) by cooling a single cryoloop (Lomber et al., 1999) placed in the lunate sulcus (loop #2 in Fig. 1a), repeated our original measurements, and then allowed cortex to rewarm before obtaining a third set of measurements.

**Figure 1.**
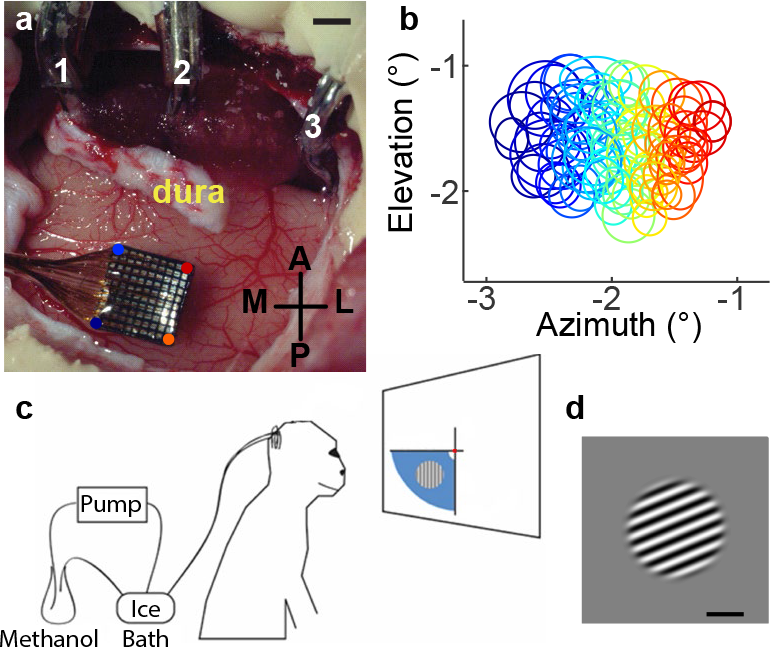
Methods. **a**, Local field potentials (LFP) and muti-unit activity (MUA) were recorded using 10×10 “Utah” multi-electrode arrays (MEA) implanted in the operculum of V1. Also shown are the top stems of the 3 cryoloops (numbered) implanted in the lunate sulcus (the actual loops are not visible as they are buried within the sulcus). **b**, Receptive fields of all electrodes on the MEA. **c**, Experimental apparatus for cortical inactivation. **d**, Visual stimulus. Scale bars, 2 mm (a), 1° of visual angle (d).

Inactivating feedback had a large effect on the overall amplitude of the LFP (Fig. 2). During control conditions, the VEP, which is the LFP aligned to stimulus onset and averaged across trials, began with a characteristic transient downward deflection approximately 40 ms after stimulus onset (‘on’ response), settling into a strong gamma rhythm during the sustained activity, and ending with a large transient at stimulus offset (‘off’ response). (The gamma rhythms are not apparent in Fig. 2 due to averaging across trials.) When feedback was inactivated, the VEP modulation was attenuated (Fig. 2a-c), with particularly large changes to the secondary phases of the ‘on’ and ‘off’ responses that occurred approximate 80-100 ms after the stimulus change. The VEP amplitude changes were clearly visible on single trials (Fig. 2d) even though spike rates (Fig. 2e) were either unchanged, as in this example, or even increased, as they were on average (see Fig. 7a). The most consistent effect of feedback inactivation was a strong reduction in the later component of the ‘on’ response, with other changes varying somewhat across electrodes and between animals (Fig. 2f,g).

**Figure 2.**
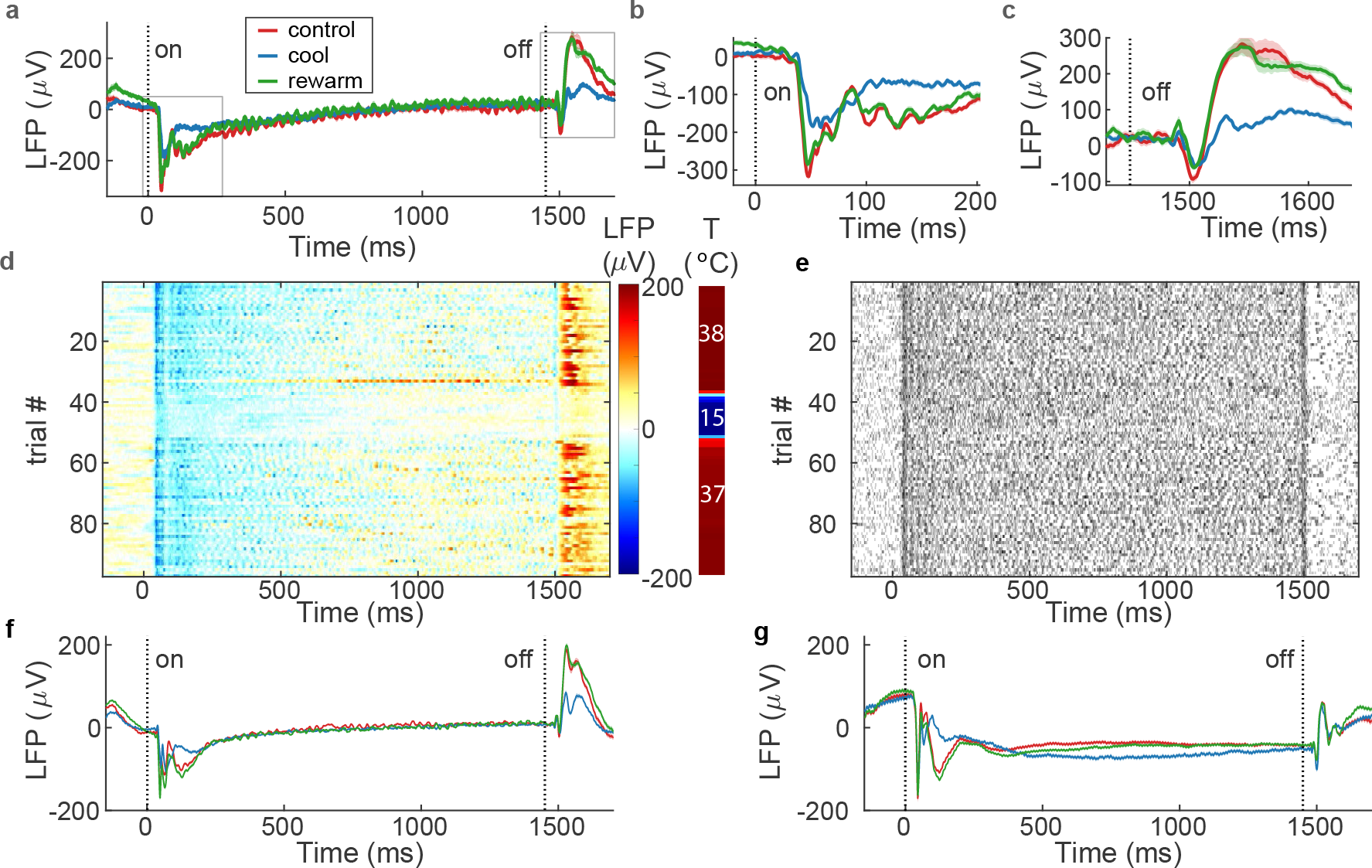
The LFP is attenuated by inactivation of feedback. Data in **a-e** are derived from the LFP recorded from one electrode in response to a large, preferred-orientation grating. Throughout this and other figures, colors indicate data recorded during control (red), cooling (blue), and after rewarming (green). **a**, Visually evoked potential for a stationary grating presented for ~1.5 s. **b**,**c**, Same data as (a), but at an expanded time scale to show details of the ‘on’ and ‘off’ responses more clearly. Shaded bands indicate ± s.e.m. (often smaller than line width). **d**, VEP for individual trials (pref. orientation). **e**, Spike rasters for the same trials shown in panel (d). **f**,**g**, Mean LFP over all electrodes for monkeys U and A, respectively.

One possible concern is that the VEPs we measured in V1 are largely the result of signals generated in V2 and propagated to V1 through volume conduction (Kajikawa and Schroeder, 2011). Insofar as this is the case, it would not be at all surprising to see a dramatic effect of inactivation on the LFP, since cooling directly suppresses synaptic activity (Lomber et al., 1999). To ascertain whether volume conduction was contaminating our VEP recordings, we conducted control experiments in which we systematically moved the visual stimulus to different locations in the visual field, thus producing differing degrees of overlap between the grating and the receptive fields of the neurons on the various electrodes on the array (Fig. 3a). Moving the visual stimulus even half a degree outside of the receptive field of the neurons on a given electrode greatly reduced the amplitude of the VEP (Fig. 3b,c) to the point where it was negligible by one degree of displacement (Fig 3d). Moreover, this effect occurred regardless of the direction in visual space that the grating was displaced—if physical distance from V2 were important, we would have expected to see a greater degree of volume conducted contamination for gratings near the vertical meridian, corresponding to the receptive fields of V2 neurons that are physically closest to V1. We thus conclude that the VEP measured in our experiments largely reflects local activity in V 1.

**Figure 3.**
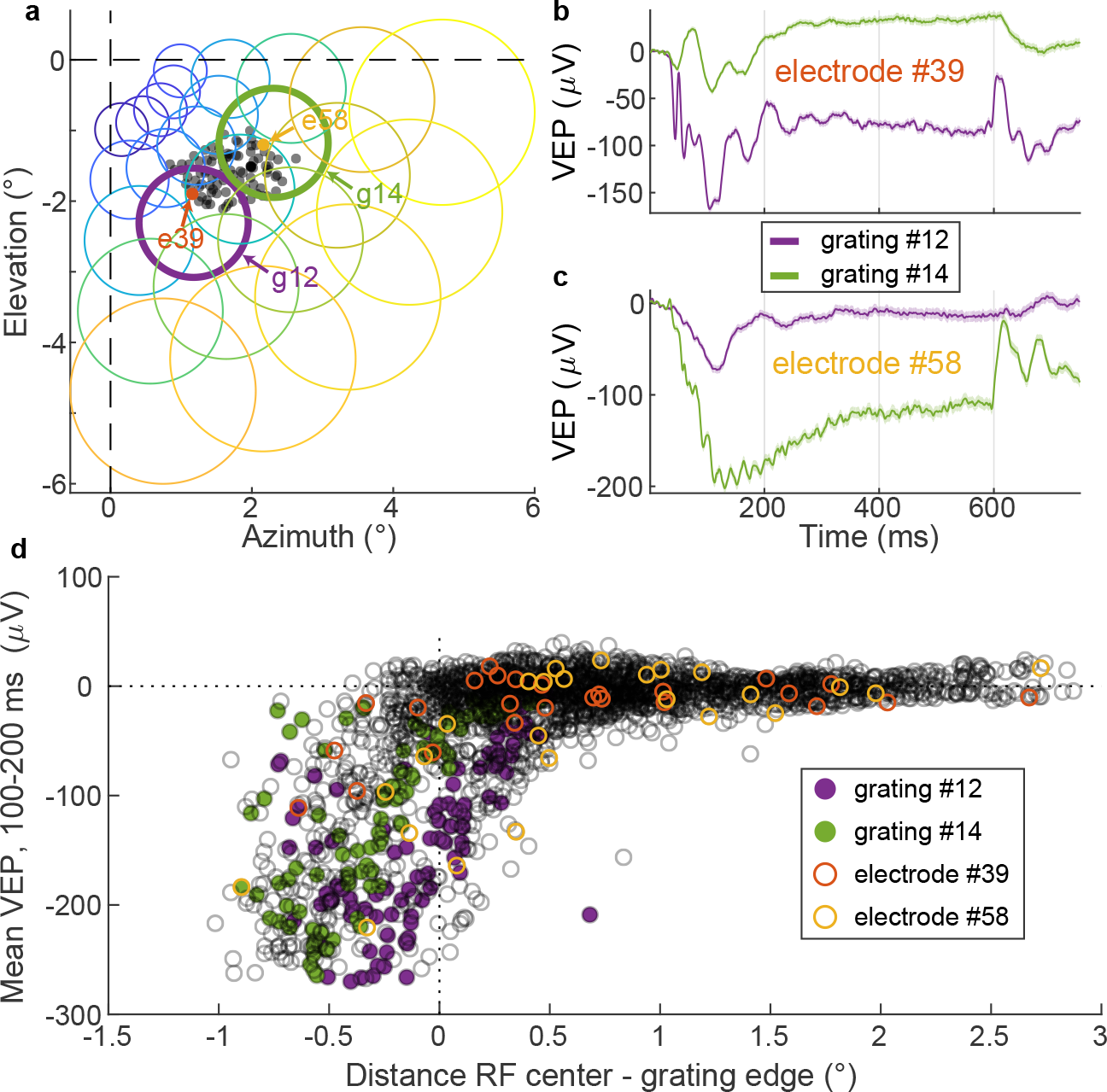
The VEP is a local phenomenon. **a**, Black dots represent receptive field (RF) centers for each recording electrode for monkey U. The orange and yellow arrows indicate two recording electrodes (e39 and e58, respectively), from which the VEPs shown in panels (b) and (c) were recorded. Each colored circle shows the outline of 1 of 25 different gratings presented in separate experiments. The purple and green arrows point to the two different gratings (g12 and g14, respectively) that evoked the responses shown in panels (b) and (c). **b,c**, The VEP for the two different recording electrodes highlighted in (a). Green and purple lines indicate the response evoked by the presentation of the grating outlined by green (g12) or purple (g14) in (a). **d**, Magnitude of the downward deflection of the VEP for all 25 grating locations and all 96 electrodes as a function of the distance between grating edge and the RF center. Data from gratings #12 and #14 are highlighted, as are the example electrodes, #39 and #58. Negative distance values indicate that the RF was inside the grating.

In addition to affecting VEP amplitude, inactivating feedback also affected LFP oscillations on many of the electrodes—an example from a single electrode is shown in Fig.4. A strong reduction in visually evoked gamma oscillations was obvious, even on individual trials (Fig. 4a-c). Because the gamma oscillations varied slightly in phase from trial-to-trial, averaging across trials would obscure them. We thus quantified the oscillations by calculating the spectral density over time for each trial using multi-taper methods (Fig. 4d-f) prior to averaging. In order to directly compare rhythms at different frequencies for control and inactivation periods, we measured the average power at each frequency across the sustained period (250-1250 ms, Fig. 4g). For this electrode, inactivating feedback reduced power at all but the lowest frequencies, but the effect was most pronounced for the gamma “bump” that peaked at about 36 Hz (Fig. 4g). This amounted to a 16-fold reduction in peak spectral power (Fig. 4h), while other frequency bands, such as alpha, were reduced by only twofold.

**Figure 4.**
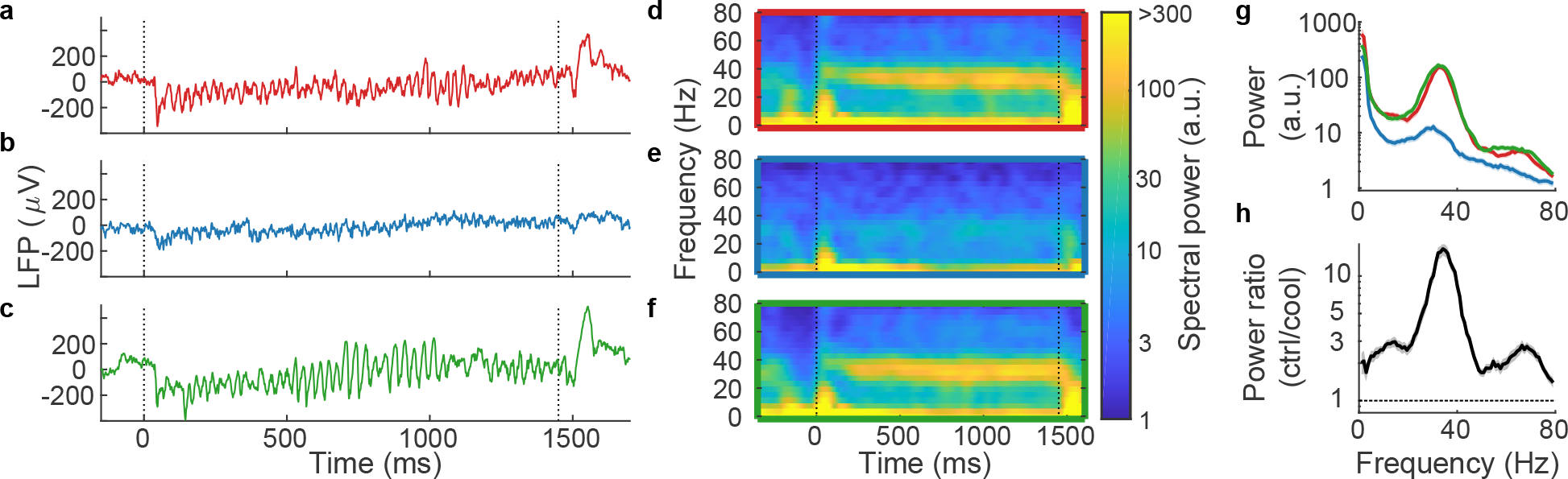
Gamma rhythms in the LFP are weakened by the inactivation of feedback; single-electrode example. **a-c**, LFP on single trials for control, cooling, and rewarm periods, respectively. **d-f**, Spectral density over time for same periods. **g**, Spectral power of LFP recorded during sustained activity. Shaded bands indicate 95% confidence intervals (Jackknife method). **h**, Power ratio of control periods (both before and after inactivation) over inactivation period. The power ratio is 16 times ±2 greater at 36 Hz without V2/V3 inactivation. The shaded area indicates bootstrapped estimates for s.e.m..

While the single-electrode example in Fig. 4 shows a pronounced effect of V2 inactivation on the gamma rhythm, the population effects were more diverse across different electrodes. As was seen for the example in Figs. 2 and 4, the mean LFP power at almost all frequencies and times was stronger when feedback was intact (Fig. 5a). The mean power ratio (control/cool) revealed three frequency peaks centered at approximately 5, 15, and 40 Hz (Fig. 5B), indicating that those frequencies were all influenced by feedback inactivation. However, it was at the peak gamma frequency that we saw the most consistent effects across electrodes: The LFP power ratio at the gamma peak was higher under control conditions at 175 of 176 sites (Fig. 5c), with a geometric mean ratio of 2.6 ±1.1. We were further motivated to focus on gamma oscillations because they are less subject to volume conduction than are lower frequencies and as will be shown below, we frequently observed strong gamma rhythms in both the LFP and the spiking activity.

**Figure 5.**
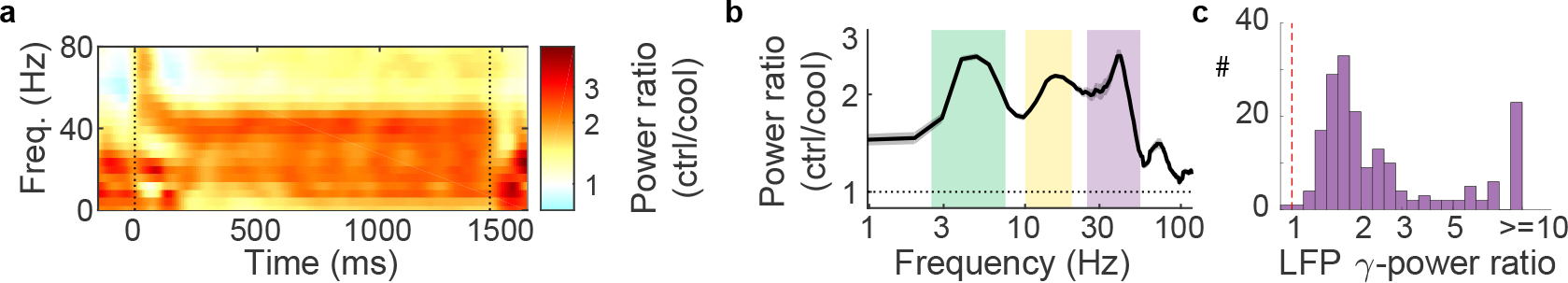
Population data for LFP oscillations. **a**, LFP spectral density ratio (ctrl/cool) averaged across all 176 electrodes. **b**, Mean sustained period (250-1250 ms after stimulus onset) power ratio plotted against frequency (log scale). Shaded gray area indicates s.e.m. **c**, Distribution of sustained period power ratio at the peak gamma frequency at all recording sites.

Interestingly, while the LFP amplitude was dramatically affected by feedback inactivation (Fig. 2), the overall spiking rates of neurons, as assessed by the multi-unit activity (MUA), were largely unaffected (Fig. 6). The MUA showed a strong stimulus driven response that looked very similar across conditions (Fig. 6a-c,e-g); however, a closer analysis revealed differences in spike timing. With feedback intact, the MUA, like the LFP, oscillated at frequencies in the gamma range (Fig. 6d) as has been previously reported (Eckhorn et al., 1988; Livingstone, 1996). But MUA gamma oscillations were completely and reversibly abolished during V2 inactivation (Fig. 6h-l), despite only minor changes in overall spike rate (Fig. 6b). This pattern was made clearer by examining the spiking autocorrelation functions (Fig. 6d), which revealed that, during control conditions, spikes were 15% ±2 more likely to occur at the peak of the gamma cycle, whereas during feedback inactivation, there was no such modulation (0.2% ±3).

**Figure 6.**
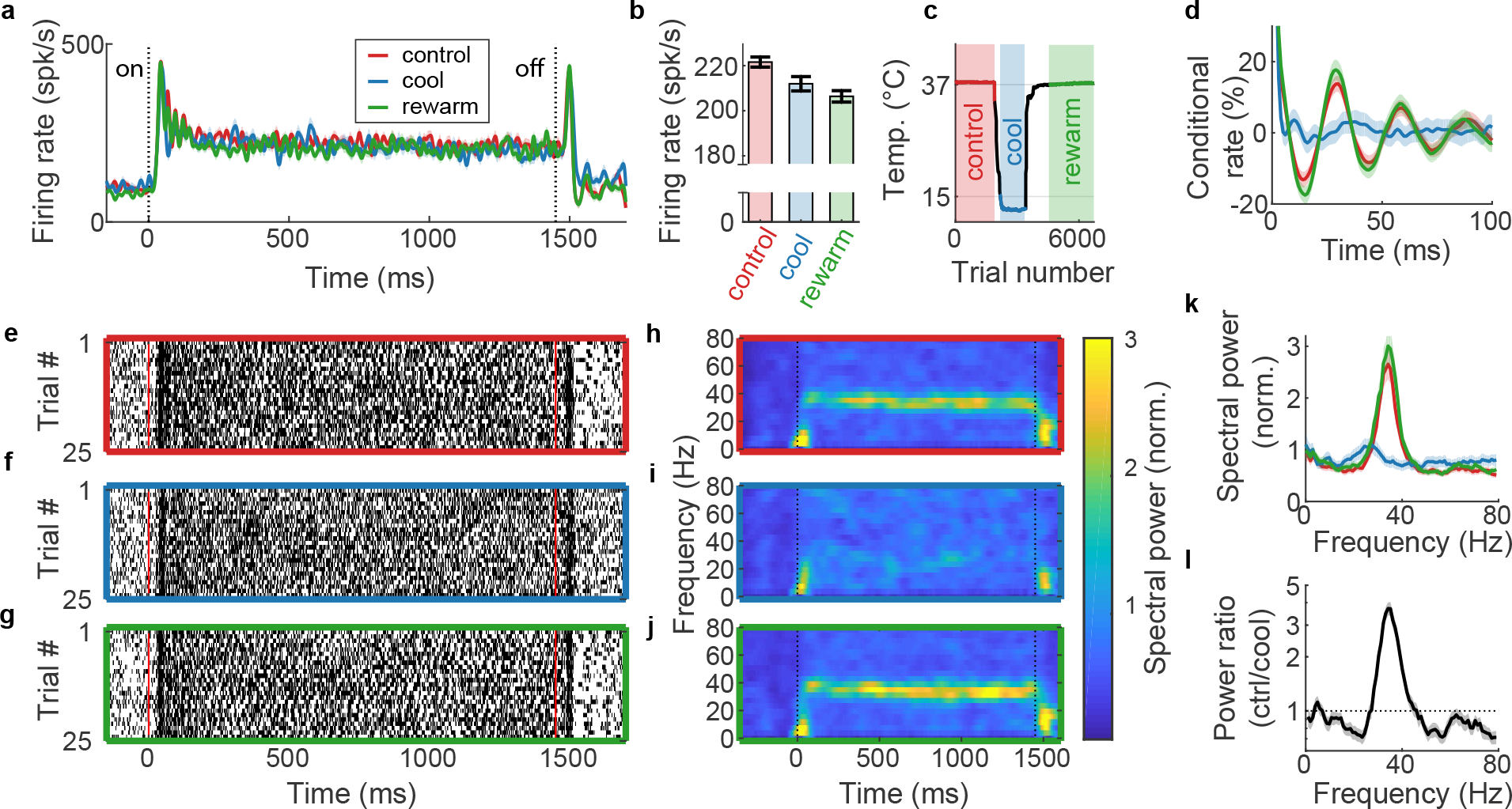
Gamma-band power of MUA disappears in the absence of V2 feedback. All spike data are derived from MUA recorded from one example electrode when a large, preferred orientation (135°) grating was displayed. **a**, Peristimulus firing rates evoked by a stationary grating. **b**, Spike rate during sustained period. Error bars indicate s.e.m. **c**, Temperature at cooling loop (near V2/V3) over the duration of the experiment. **d**, Autocorrelation of MUA during sustained activity. A conditional rate of 20% (at *e.g.* 30 ms) means that spikes were 20% more likely to be recorded 30 ms after reference spike occurrence, relative to the average rate. **e-g**, Raster plots for 25 random trials for control, cooling, and after rewarming periods, respectively. **h-j**, Spectral density, derived from MUA spiking, for control, cooling, and after rewarming periods, respectively. The power is normalized; a Poisson process would have a power of one. **k**, Spectral power of frequency of spikes recorded during sustained activity. Shaded bands indicate 95% confidence intervals (Jackknife method). **l**, Ratio of power recorded during control conditions over power during feedback inactivation. The power ratio is 3.7 times ±0.3 higher at 36 Hz with feedback intact. Shaded areas are bootstrap estimates for s.e.m.

The patterns in MUA revealed at individual electrodes (Fig. 6) were largely consistent across the 176 electrodes recorded in two animals (Fig. 7). Overall, spike rates increased by an average of 5 (±1) spk/s during feedback inactivation (Fig. 7a). Recall that we used large (~5° diameter) gratings that invaded the suppressive surrounds of V1 neurons. The increase in spike rate during cooling is therefore consistent with previous reports that feedback inactivation reduces surround suppression (Nassi et al., 2013, 2014; Nurminen et al., 2018). Despite this increase in spike rate, the periodicity in spiking decreased across the vast majority of sites, whether measured as a change in gamma power (Fig. 7b) or as a change in the conditional probability of spiking (CPS, Fig. 7c). This was also reflected in the mean power ratios of the MUA activity, which showed that the effect of feedback inactivation was strongest in the gamma band (Fig. 7d,f). And these analyses considerably underestimate the effect of feedback inactivation on spike timing, because many electrodes showed very little MUA gamma power under control conditions. When we restricted our analyzes to the subset of electrodes (n=64) for which there was significant gamma during control (p<0.05, Jacknife test), the effects were stronger (Fig. 7e,g) and the distribution of power ratios shifted considerably to the right (Fig. 7h).

**Figure 7.**
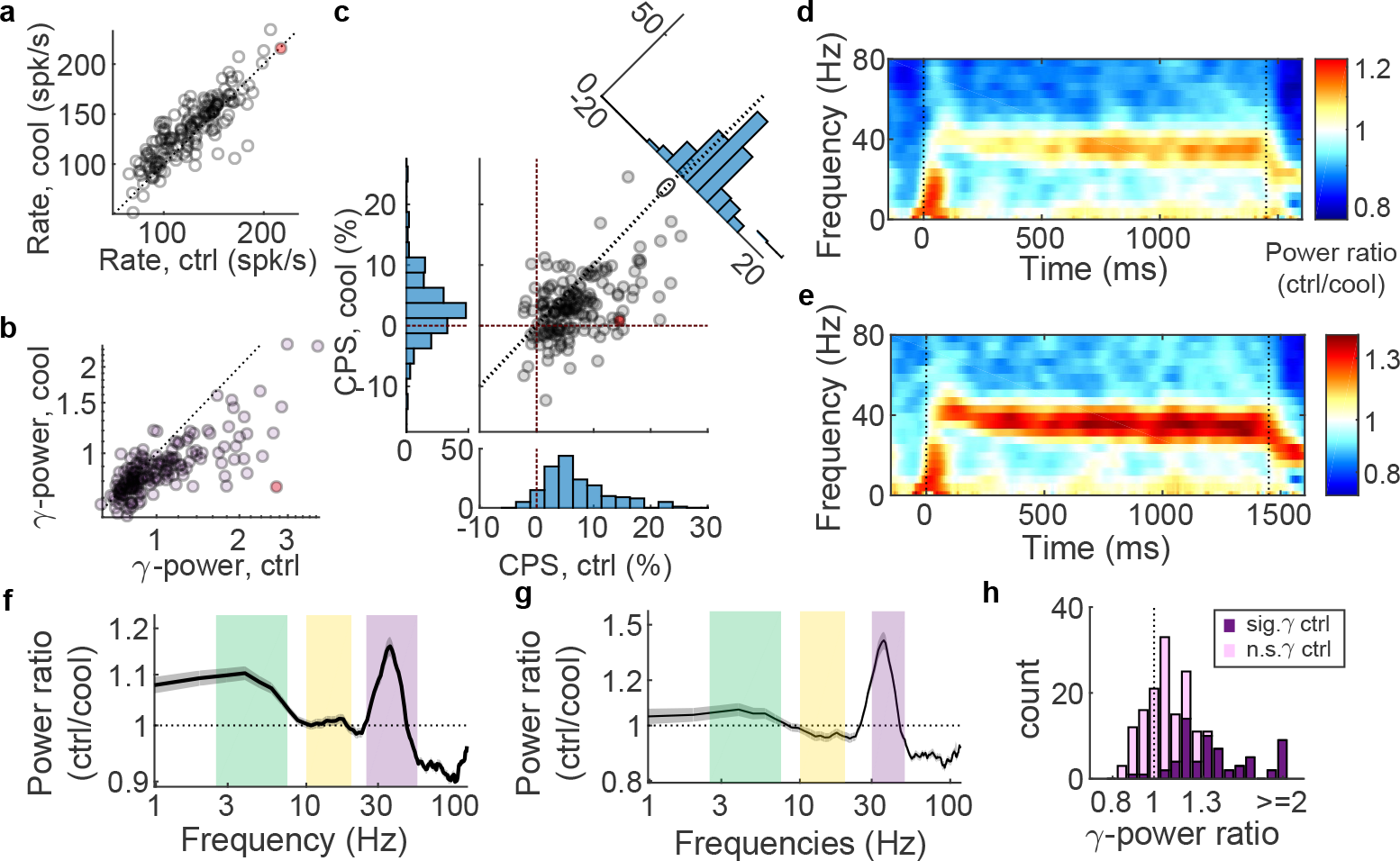
Population data for MUA shows dependency of oscillations on feedback. **a**, Spike rate for all sites (n = 176) during controls and during feedback inactivation. **b**, Power during control and feedback inactivation during the sustained period at peak gamma frequency (MUA). The mean decreased from 1.11 (±0.04) during controls to 0.89 (±0.02) during inactivation (p< 10^−10^, paired t test). **c**, Conditional probability of spiking (CPS) from autocorrelation at the time point based on the peak of the gamma oscillation. The mean decreased from 7.3% (±0.5%) during controls to 2.9% (±0.4%) during inactivation (p< 10^−10^, paired t test). The red dot in (a-c) indicates the single-electrode data from Fig. 6. **d**, Mean power ratio (power during control conditions divided by power during feedback inactivation) over time and for different frequencies. **e**, Same as panel (d) but for the subset of electrodes that exhibited significant MUA gamma (p<0.05) during the control period. **f**, Mean sustained response power ratio plotted against frequency (log scale) for all electrodes. Shaded gray area indicates s.e.m. **g**, Same as panel (f) but for the subset of electrodes that exhibited significant MUA gamma (p<0.05) during the control period. **h**, Distribution of sustained period power ratio at gamma frequency peak. Light purple indicates MUA without significant gamma oscillations during the control period, dark purple shows MUA with significant gamma oscillations.

To explore possible underlying mechanisms, we made use of a previously published conductance-based model that specifically included feedback from extrastriate cortex in order to explain the dependence of the peak frequency of V1 gamma rhythms on the size of the visual stimulus (Jia et al., 2013a; Kang et al., 2010). The model consists of two V1 populations, one excitatory (E) and one inhibitory (I), that are connected both to each other and to themselves (recurrence), typical of models that produce “pyramidal-interneuron network gamma,” or PING (Börgers and Kopell, 2003; Tiesinga and Sejnowski, 2009; Whittington et al., 2000). Importantly, the model of (Kang et al., 2010) adds a projection from the V1 E neurons to a V2 population that integrates the V1 activity and then projects back to both the E and I populations in V1 (Fig. 8a). By slightly modifying model parameters—mainly strengthening the influence of feedback and weakening that of local recurrent connections—we were able to reproduce our main experimental finding of the dependence of gamma on cortico-cortical feedback (Fig. 8b-e). In our modified version of the model, the V1 gamma rhythm is caused by both local excitatory-inhibitory interactions and cortico-cortical feedback; neither alone can produce the rhythm. The conductance patterns in the modified model exhibit striking similarities to experimental results (where firing rates were measured), including increased spike rates when feedback is removed (Nassi et al. (2013, 2014) and present results), a consistent lag of V2 gamma oscillations by approximately 3-5 ms (Jia et al., 2013b), and trial-by-trial fluctuations in gamma amplitude and frequency that covary in V1 and V2 (Frien et al., 1994; Roberts et al., 2013).

**Figure 8.**
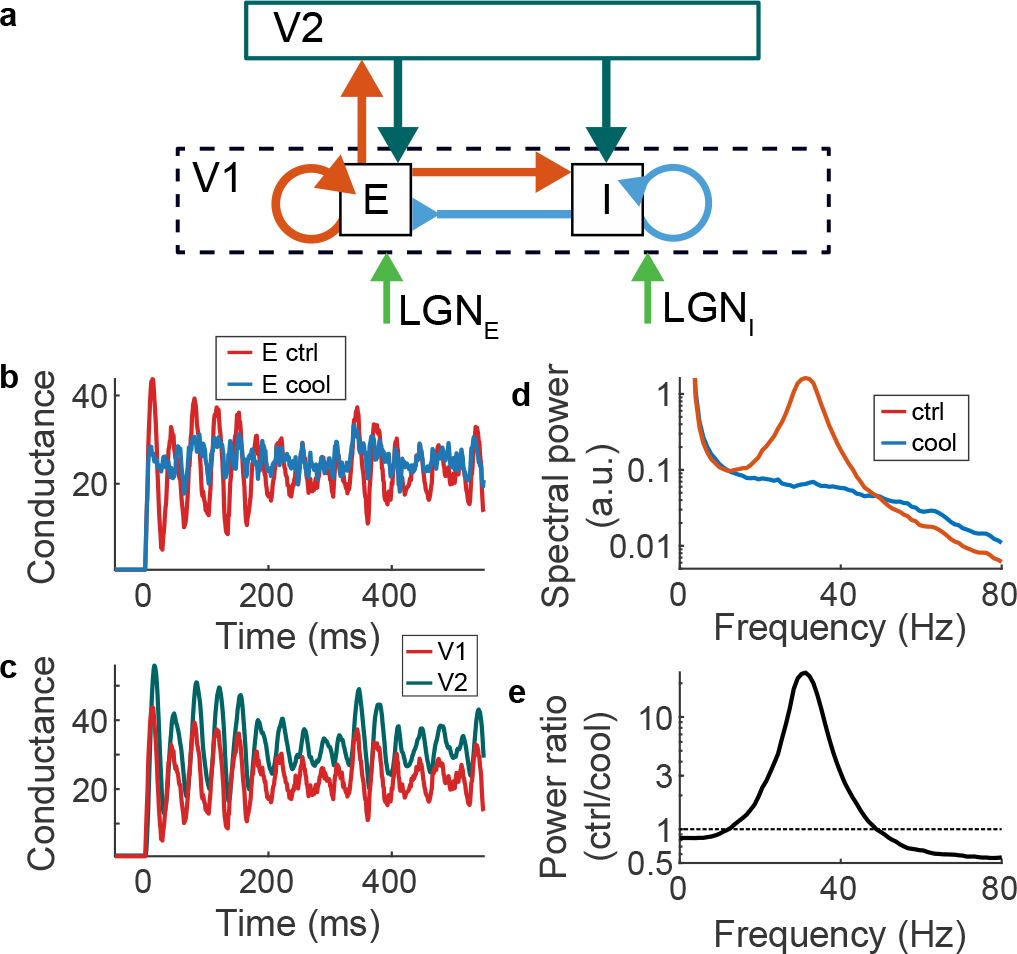
Feedback is required for gamma oscillations in the model. **a**, Implementation of a conductance based model (Kang et al., 2010) with three neuronal populations: excitatory (E) and inhibitory (I) V1 neurons as well as V2 neurons with long range projections to V1. Both E neurons and I neurons in V1 receive LGN input. E projects onto itself and I locally as well as to V2. I only directly inhibits itself and local E, but not V2. V2 projects to both E and I. **b**, Modelled conductance response to stimulus onset with full model (red, ctrl) and when eliminating V2 feedback connections (blue, cool). The recurrent loop from V1 to V2 is necessary for the gamma oscillation for the parameters we chose. **c**, V2 neuronal conductance (teal) follows the excitatory population in V1 (red). **d**, Spectral power of the E population during the sustained period. **e**, Power ratio during sustained period (ctrl/cool).

## 3. Discussion

Our experiments have revealed a surprisingly strong influence of cortico-cortical feedback from V2 on both the LFP and spike timing in V1, even when spike rates in V1 were only weakly affected. Perhaps the most striking finding was the complete disappearance of stimulus-evoked gamma in the MUA with corresponding changes in the autocorrelation of V1 spiking activity.

We are confident that the effects we observed were due to feedback inactivation and not to any direct cooling of V1. In previous studies (Nassi et al., 2013), we made direct measurements of cortical temperature in V1 when we inactivated V2/V3 with three adjacent cryoloops in the lunate sulcus, each cooled to between 1 and 5°C. Under these conditions, direct cooling in V1 amounted to <2°C. In the present study, we cooled only a single loop to between 14 and 15°C (Fig. 2d), which should make direct cooling of our V1 recording sites negligible. In addition, in our previous control experiments we found that direct cooling only produced decreases in spike rates (see Nassi et al., 2013, Fig. 9), whereas here we observed spike-rate increases, consistent with a reduction in surround suppression reported previously (Nassi et al., 2013, 2014). Further evidence against direct cooling is the fact that the distance of the V1 recording sites from the lunate sulcus, where the cryoloop resided, showed no consistent relationship with the strength of the effects of cooling on cortical rhythms.

The current consensus is that gamma rhythms are generated locally, within a cortical area, through recurrent loops involving fast spiking inhibitory interneurons (Börgers and Kopell, 2003; Buzsáki and Wang, 2012; Chariker et al., 2018; Headley and Paré, 2017; Tiesinga and Sejnowski, 2009). The fact that they can be produced in reduced preparations, such as cortical slices (Whittington et al., 1995), makes a strong argument for the sufficiency of local circuits. But our finding that gamma rhythms in V1 depend on feedback from V2 supports the idea that gamma oscillations may also involve long-range connections between cortical areas. Another possibility, however, is that visually evoked gamma is a purely local rhythm, but is subject to control by remote sources. If, for example, feedback targeted interneurons that were critical for the oscillation, then even relatively subtle modulatory effects, such as changes in the decay time of inhibition (Kopell et al., 2010), could lead to large changes in the rhythm-generating capacity of the local circuit.

Related to this second possibility and to previous findings that the peak frequency of gamma rhythms are modulated by stimulus size (Gieselmann and Thiele, 2008; Henrie and Shapley, 2005; Ray and Maunsell, 2010), we note two similarities between the effects of inactivating feedback to V1 in the monkey and local inactivation of somatostatin-expressing inhibitory interneurons (SOMs) in V1 of mice. In monkeys, V2/V3 inactivation produced a reduction in surround suppression in V1 (Nassi et al., 2013, 2014; Nurminen et al., 2018) very similar to that seen in mouse V1 when SOMs were optogenetically silenced (Adesnik et al., 2012). And our current finding of an attenuation of gamma rhythms with feedback inactivation echoes a more recent study (Veit et al., 2017), which also found a reduction in visually evoked gamma oscillations in mouse V1 when SOMs were suppressed. Although evidence for a direct projection from V2 feedback neurons that selectively targets SOMs is lacking (Gonchar and Burkhalter, 2003), these similarities indicate that one of the mechanisms by which corticocortical feedback influences local computations is through an indirect interaction with local SOM microcircuits.

A possible SOM-feedback interaction is further reinforced by the anatomical evidence that the layer 1 apical dendrites of pyramidal neurons (whose cell bodies reside in layers 2, 3 or 5) are a major target of both feedback from V2 (Anderson and Martin, 2009; Johnson and Burkhalter, 1996, 1997; Rockland and Virga, 1989) and of SOMs (Kapfer et al., 2007; Ma et al., 2006; McGarry et al., 2010; Silberberg and Markram, 2007; Wang et al., 2004). Our present and previous (Gomez-Laberge et al., 2016) findings can be plausibly accounted for by this circuitry when we consider that layer 5 pyramidal neurons can act as coincidence detectors in which input to the apical dendrites, when paired within a narrow time window with input to the soma, can produce bursts of action potentials, as opposed to single spikes when no distal input is supplied (Larkum et al., 1999, 2001). There is strong evidence that a subtype of SOMs, the so-called Martinotti cells, directly gate the excitability of pyramidal neuron apical dendrites (Cichon and Gan, 2015; Gentet et al., 2012; Murayama et al., 2009; Silberberg and Markram, 2007; Takahashi et al., 2016), consistent with the finding that inhibiting SOMs produced an increase in burst firing of pyramidal cells (Gentet et al., 2012). A switch from burst mode to single-spiking, whether mediated by SOMs or by a direct effect of feedback on the apical dendrites, could account for the reduced spiking variability we observed with feedback inactivation (Gomez-Laberge et al., 2016), could at least partially explain feedback’s powerful influence over the volley of recurrent activity that follows the initial response in the VEP (Fig. 2), and could also produce strong effects on local rhythms.

Finally, several groups have argued that cortical rhythms in different frequency bands are characteristic of differences in the direction of information flow within cortical hierarchies, specifically linking gamma rhythms to feedforward processing and slower rhythms (in the alpha or beta range) to feedback (Bastos et al., 2015; van Kerkoerle et al., 2014). Our data argue against the strongest form of this framework, since by reducing feedback from V2 to V1 we observed profound reductions in gamma oscillations even as feedforward drive was unchanged. However, our results do not rule out the possibility that the direction of information transfer within different frequency bands, as measured by techniques like Granger causality, is indeed in the proposed directions. Whether gamma rhythms play a critical role in cortical computation, or whether they are an epiphenomenal signature of particular circuit motifs, remains a fascinating and open question.

## Acknowledgment

We thank Camille Gómez-Laberge and Vladimir Berezovskii for technical help and discussions as well as Gabriel Kreiman, John Maunsell, and Nancy Kopell for comments on the manuscript. This work was supported by F32 EY025523 (TSH), R01 EY011379 (RTB), and P30 EY012196 (core grant).

## Author Contributions

T.S.H and R.T.B. conceived experiments. S.G.L fabricated the cryoloops and, along with T.S.H. and R.T.B., implanted them in the monkeys. T.S.H. and S.R. performed experiments. T.S.H. analyzed data and model. T.S.H. and R.T.B. wrote the paper. T.S.H. and R.T.B. acquired funding.

## 4. Online Methods

### 4.1. Experimental Model and Subject Details

All experiments were approved by the Harvard Medical Area Standing Committee on Animals and were in accordance with the National Institutes of Health Guide for Care and Use of Laboratory Animals.

### 4.2. Method Summary

The experimental details have been described previously (Nassi et al., 2013; Trott and Born, 2015). Briefly, two male rhesus monkeys (*Macaca mulatta*), A and U, were implanted with a head post, a multi-electrode array (MEA) over the V1 operculum, and three custom made cryoloops (Lomber et al., 1999) into the lunate sulcus in an aseptic surgery under isoflurane anesthesia. The electrodes on the MEA were 1 mm long and spaced 400 *μm* apart in a 10×10 grid. We took photographs during the surgery and used them to estimate the distance between the electrodes on the electrode array and the location where the loop enters the lunate sulcus. The animals learned to maintain fixation within a 1.5 to 2° window for liquid juice rewards while large circular sinusoidal grating patches appeared on a computer screen in front of them.

### 4.3. Stimuli

All stimuli were created in MATLAB (Natick, MA) with custom computer code using Psychtoolbox (Kleiner et al., 2007). Gratings were displayed 75 cm in front of the animals’ eyes on a 40×30 cm CRT screen with a refresh rate of 100 Hz. Gratings were displayed for 1450 ms centered over the aggregate RFs recorded from the MEAs. All recorded RFs were completely covered by the 3.6° (monkey U) and 5° (monkey A) diameter gratings. We displayed 8 orientations (0 to 157.5°) at 100% contrast. The spatial frequencies of the gratings were 4 cycles/° (monkey U) and 2 cycles/° (monkey A); gratings remained stationary for the duration of each trial.

### 4.4 Electrophysiology

We recorded the raw voltage on all electrodes continuously (digitized at 30 kHz) during the experiments with the Blackrock Microsystems Cerebus system. Offline, we extracted the LFP by low pass filtering the raw signal (<200 Hz) and aligning it to stimulus onsets. To record the action potentials of nearby neurons — the MUA — we band pass filtered the raw signal (>250 Hz and <5 kHz) and extracted events when the voltage exceeded −2.4 times the standard deviation of the signal. We visually inspected the mean voltage around the threshold crossing and confirmed that the signal followed the typical action potential waveform. To measure orientation tuning, for each electrode we fitted von Mises functions to the mean MUA firing rate for each orientation and defined the preferred orientation as the stimulus orientation that was closest to the peak of the von Mises function. We also continuously monitored the temperature on the cryoloops using a thermocouple embedded in the loop (Lomber et al., 1999). Each experimental session consisted of three periods: 1) control trials, prior to cooling, 2) cool trials, during which the middle cryoloop was cooled to between 14 and 15°C, and 3) rewarm trials, which began approximately 20 min after the cryoloop temperature returned to above 36°C. We did not find differences in any of the spectral density metrics between the control and rewarm periods, so we combined data from both periods for population analyses.

**Table 1.**
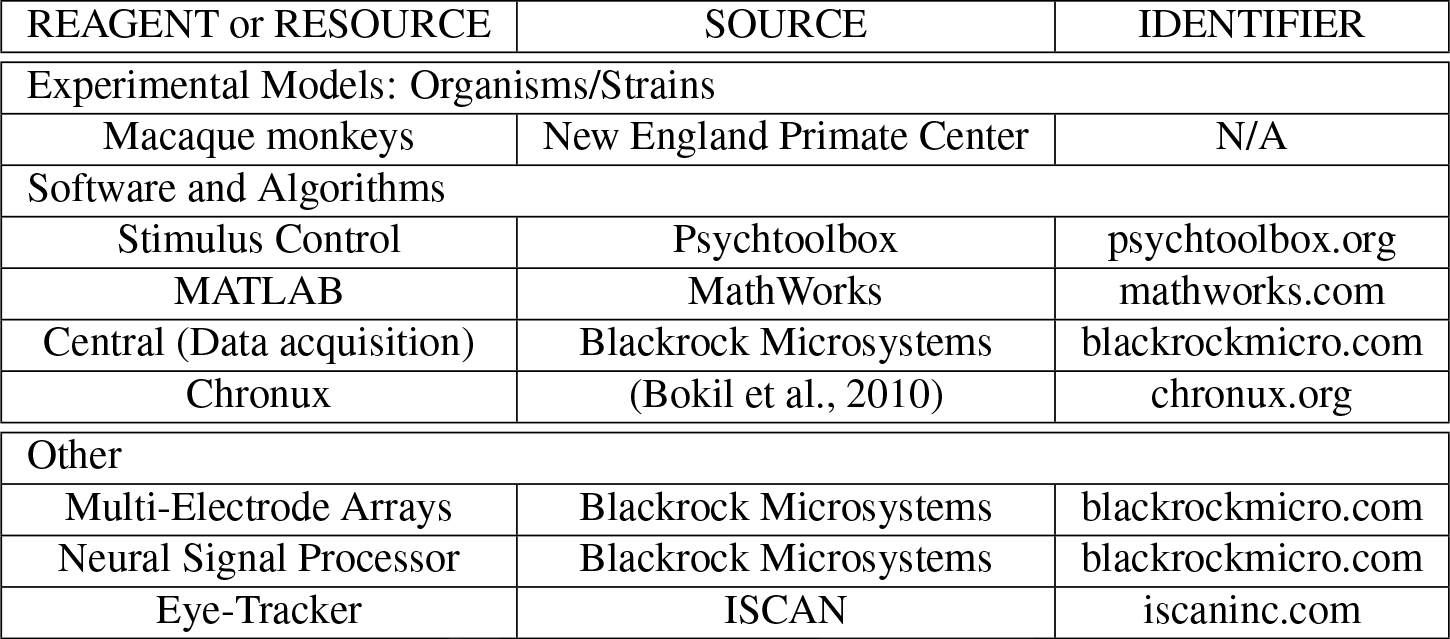
KEY RESOURCES TABLE

| REAGENT or RESOURCE | SOURCE | IDENTIFIER |
| --- | --- | --- |
| Experimental Models: Organisms/Strains |  |  |
| Macaque monkeys | New England Primate Center | N/A |
| Software and Algorithms |  |  |
| Stimulus Control | Psychtoolbox | psychtoolbox.org |
| MATLAB | MathWorks | mathworks.com |
| Central (Data acquisition) | Blackrock Microsystems | blackrockmicro.com |
| Chronux | (Bokil et al., 2010) | chronux.org |
| Other |  |  |
| Multi-Electrode Arrays | Blackrock Microsystems | blackrockmicro.com |
| Neural Signal Processor | Blackrock Microsystems | blackrockmicro.com |
| Eye-Tracker | ISCAN | iscaninc.com |

### 4.5. Spectral analysis and autocorrelation

We estimated the spectral power around the grating presentations (100 ms before onset to 100 ms after stimulus offset) using the multi-taper method from the Chronux toolbox (Bokil et al., 2010) (parameters: time-bandwidth product TW = 2, time window T = 0.2 s, number tapers k = 1) for the LFP, the MUA, and the modelled conductances for each trial, then averaged all trials for a given condition. To estimate the spectral power during the sustained activity, from 250 to 1250 ms after stimulus onset, we used multi-taper with a time-bandwidth product TW = 3 and the number of tapers k = 5. For examples displayed in Fig. 4g and 6k, we estimated the 95% confidence intervals with a Jackknife procedure (Miller, 1974). For the s.e.m. in Fig. 4h and 6l, we performed bootstrap analysis by resampling (with replacement) the trials for both control and inactivation periods 1000 times. The MUA autocorrelation was determined with a modified version of MATLAB’s autocorrelation function (autocorr). To estimate the conditional spiking probability at the gamma peak, we measured the conditional rate at the time point expected from the peak gamma frequency (monkey A, t = 1/44 Hz = 23 ms; monkey U, t = 1/35 Hz = 29 ms).

### 4.6. Population Statistics

We recorded signals from 96 electrodes in each animal. Some of these electrodes had strong line noise artifacts, so we excluded electrodes (n = 16) where the spectral power at the first harmonic (120 Hz) exceeded 1.2 times the mean power between 115 to 117 Hz and 123 to 125 Hz. To estimate the effect of feedback inactivation on frequency over time (Fig. 3A,B), we first calculated the spectral density over time for each condition and then divided the spectral density during the control periods by the spectral density during inactivation (see above). We then took the mean for each frequency/time value for all channels and displayed the result in log scale. Similarly, we calculated the population power ratio over the sustained activity by dividing the spectral power during the controls by the spectral power during inactivation. The two monkeys had slightly different peak gamma band frequencies (monkey A, 44 Hz; monkey U, 35 Hz), so for the comparison of changes in gamma power, we measured power at the respective peak frequencies.

### 4.7. Modelling Details

We explored feedback dependent gamma rhythms in V1 using a previously published model (Jia et al., 2013b; Kang et al., 2010), that simulates the conductances of two V1 populations — one excitatory (E), one inhibitory (I) — and one V2 population (V2). The rate-of-change of the conductances was estimated as:

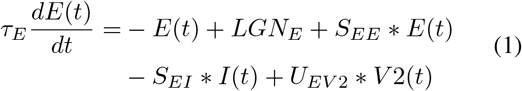

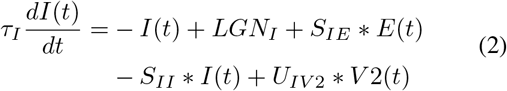

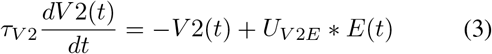

Equations 1, 2, and 3 are 2.1-2.3 from Kang et al. (2010), integrated over space. We calculated the conductances for 1700 ms, starting at t=−200 with a very small LGN input that became strong at t=0 ms and remained so until t=1450 ms, at which time the LGN input became weak again. We numerically calculated the change of E, I, and V2 at each millisecond according to the above equations and added the results to the conductances. We repeated 100 stimulus presentations to estimate spectral power density (same settings as above). See Table 2 for details.

**Table 2.** Model parameters

|  |  |  |
| --- | --- | --- |
| $E(t)$ | $E(-400) = 0$ | Conductance rates of excitatory population |
| $I(t)$ | $I(-400) = 0$ | Conductance rates of inhibitory population |
| $V2(t)$ | $V2(-400) = 0$ | Conductance rates of V2 population |
| $\tau_E$ | $4ms$ | Excitatory synaptic time constant |
| $\tau_I$ | $6ms$ | Inhibitory synaptic time constant |
| $\tau_{V2}$ | $4ms$ | V2 synaptic time constant |
| $LGN_E$ | $Poission(\lambda = 20)$ | Ex. LGN input (0-1450 ms): Poission noise added, independently of $LGN_I$ |
| $LGN_I$ | $Poission(\lambda = 5)$ | Inh. LGN input (0-1450 ms): Poission noise added |
| $S_{EE}$ | 1.3 | Connection strength onto excitatory from excitatory |
| $S_{EI}$ | 1.6 | Connection strength onto excitatory from inhibitory |
| $S_{IE}$ | 1.6 | Connection strength onto inhibitory from excitatory |
| $S_{II}$ | 1.6 | Connection strength onto inhibitory from inhibitory |
| $U_{V2E}$ | 1.4 | Connection strength onto V2 from excitatory |
| $U_{EV2}$ | 1 | Connection strength onto excitatory from V2 |
| $U_{IV2}$ | 1.7 | Connection strength onto inhibitory from V2 |

